# *In vitro* broad-spectrum antiviral activity of MIT-001, a mitochondria-targeted reactive oxygen species scavenger, against severe acute respiratory syndrome coronavirus 2 and multiple zoonotic viruses

**DOI:** 10.1101/2023.07.06.547945

**Authors:** Taehun Lim, Shivani Rajoriya, Bohyeon Kim, Augustine Natasha, Hyeonjoo Im, Hyun Soo Shim, Junsang Yoo, Jong Woo Kim, Eun-Woo Lee, Hye Jin Shin, Soon Ha Kim, Won-Keun Kim

## Abstract

The COVID-19 pandemic caused by SARS-CoV-2 becomes a serious threat to global health and requires the development of effective antiviral therapies. Current therapies that target viral proteins have limited efficacy with side effects. In this study, we investigated the antiviral activity of MIT-001, a small molecule reactive oxygen species (ROS) scavenger targeting mitochondria, against SARS-CoV-2 and other zoonotic viruses *in vitro*. The antiviral activity of MIT-001 was quantified by RT-qPCR and plaque assay. We also evaluated the functional analysis of MIT-001 by JC-1 staining to measure mitochondrial depolarization, total RNA sequencing to investigate gene expression changes, and immunoblot to quantify protein expression levels. The results showed that MIT-001 effectively inhibited the replication of B.1.617.2 and BA.1 strains, Zika virus, Seoul virus, and Vaccinia virus. Treatment with MIT-001 restored the expression of heme oxygenase-1 (*HMOX1*) and NAD(P)H: quinone oxidoreductase 1 (*NqO1*) genes, anti-oxidant enzymes reduced by SARS-CoV-2, to normal levels. The presence of MIT-001 also alleviated mitochondrial depolarization caused by SARS-CoV-2 infection. These findings highlight the potential of MIT-001 as a broad-spectrum antiviral compound that targets for zoonotic RNA and DNA viruses, providing a promising therapeutic approach to combat viral infection.

## 1. Introduction

Coronavirus Disease 2019 (COVID-19^1^) began in Wuhan, Hubei province, China, in 2019. On 30 January 2020, the World Health Organization (WHO) declared a global emergency over the novel coronavirus outbreak (World Health Organization, 2020a). Subsequently, on 11 March 2020, the WHO declared COVID-19 a pandemic (World Health Organization, 2020b). According to the WHO Health Emergency Dashboard, by May 2023, approximately 820 million confirmed cases and 1 million deaths had been reported (WHO Health Emergency Dashboard). COVID-19 is an infectious disease caused by the severe acute respiratory syndrome 2 virus (SARS-CoV-2). The outbreak of COVID-19 has severely impacted the global economy and human health.

In the quest for effective antiviral treatments against COVID-19, recent studies have focused on viral genetic components and host genetic factors as main potential targets. Molnupiravir, a nucleotide analog inducing replication errors, is an approved antiviral agent which is disrupting viral replication, whereas the combination therapy of Nirmatrelvir and Ritonavir targets the viral protease (Mpro) against SARS-CoV-2 (Saravolatz et al., 2023; Lamb, Y. N., 2022). Simultaneously, host factors regulating viral susceptibility and immune response pose a critical target for developing antiviral countermeasures. 4-Octyl itaconate (4-OI), activating the Nuclear factor erythroid-2-related factor 2 (Nrf2)-related pathway, enhanced antioxidant defenses and inhibited pro-inflammatory responses upon viral infections, holding potential for antiviral effects against SARS-CoV-2 (Olagnier et al., 2020). These multifaceted strategies demonstrate the diverse approaches being pursued to develop effective antiviral therapies against COVID-19 and multiple viral outbreaks.

Here, we introduce a potential mitochondria-targeting anti-inflammatory and anti-reactive oxygen species (ROS) agent, MIT-001 (Known as NecroX-7) against SARS-CoV-2 and multiple viruses *in vitro*. MIT-001 inhibits ROS and calcium accumulation in mitochondria, which suppresses the release of damage-associated molecules by accidental necrosis and the agent-induced antioxidant action of nuclear factor kappa-light-chain-enhancer of activated B cells (NF-кB) and inflammasome-dependent cytokines (Grootaert et al., 2016; Im et al., 2015). Currently, the antiviral activity of MIT-001 remains to be investigated.

In this study, the antiviral activity of MIT-001 was evaluated against SARS-CoV-2 *in vitro*. In addition, we examined the efficacy of MIT-001 in DNA and RNA viruses to verify its broad-spectrum antiviral activity. MIT-001 could serve as a broad-spectrum antiviral agent for the treatment of SARS-CoV-2 and variants (B.1.617.2 and BA.1 strains), Seoul virus (SEOV), Zika virus (ZIKV), and Vaccina virus (VACV).

## 2. Materials and Methods

### 2.1 Ethics

The study of multiple anti-viral reagents against SARS-CoV-2 was performed at biosafety level-3 (facilities at Hallym Clinical and Translational Science Institute, Hallym University, Chuncheon, South Korea) under guidelines and protocols in line with the institutional biosafety requirements (Hallym2020-04, 30th, Oct., 2020, Hallym University Institutional Biosafety Committee). Experiments using ZIKV, VACV, and SEOV were performed at biosafety level-2.

### 2.2. Cell lines

Vero E6 (ATCC® CRL-1596) cells were cultivated in Dulbecco’s modified Eagle’ medium (DMEM, 11995065, Gibco®, Life technologies, Europe B.V) supplemented with 10% Fetal Bovine Serum (FBS, 10082147, Gibco®), 1% 10 mM HEPES in 0.85% NaCl (17-737E, Lonza, BioWhittaker®, Walkersville, MD, USA), 1% antibiotic-antimycotic, penicillin 10 U/mL, streptomycin 100 µg/mL, and Fungizone™ (amphotericin B) 0.25 µg/mL (15240062, Gibco™). A549 lung carcinoma cells expressing human ACE2 (M08-0801, ©InvivoGen, San Diego, USA) were cultivated in DMEM (Gibco®) supplemented with 10% FBS (10082147, Gibco®), 1% 10 mM HEPES in 0.85% NaCl (17-737E, Lonza), 100 U/mL penicillin, 100 µg/mL streptomycin (15140-12, Gibco®), and 100 µg/mL Normocin™ (ant-nr-1, ©InvivoGen). Cells were maintained at 37°C with 5% CO_2_.

### 2.3. Viruses

SARS-CoV-2 B.1 (NCCP No. 43326), B.1.617.2, (NCCP No. 43406), and BA.1, (NCCP No. 43408), ZIKV (NCCP No. 43280), and VACV (NCCP No. 43281) were acquired from the National Culture Collection for Pathogens (Osong, ROK). SEOV, (Korea Bank for Pathogenic Viruses, LML-14-179).

### 2.4. In vitro infection and MIT-001 treatment

Vero E6 and hACE2-A549 cells were seeded in 6-well plates (Falcon®) and incubated overnight. Upon reaching about 80% confluence, cells were washed with phosphate buffered saline (PBS, 70011069, Gibco™) and infected with viruses at specific multiplicities of infectivity (MOI, (Vero E6 MOI = 0.01, hACE2-A549 MOI = 0.1) for 2 h. MIT-001 (C_24_H_29_N_3_O_3_S, 439.57 g/mol) treatment was then applied at varying concentrations. The infected plates were manually shaken every 15 min to efficiently distribute the inoculum. The infected cells were harvest at 24 and 48 hours post-infection (hpi).

### 2.5. RNA extraction and reverse transcription-polymerase chain reaction (RT-PCR)

Total RNA was extracted from cells using TRIzol (15596026, Ambion, Life Technologies, Carlsbad, CA, USA) according to the manufacturer’s protocol. Subsequently, the extracted RNA was subjected to reverse transcription to complementary DNA (cDNA) using a high-capacity RNA-to-cDNA kit (4387406, Thermo Fisher Scientific Baltics UAB) and a SimpliAmp Thermal Cycler (A24811, Thermo Fisher Scientific). The reverse transcription reaction was performed at 37 °C for 60 minutes, followed by a denaturation step at 95 °C for 5 minutes.

### 2.6. Real-time quantitative PCR (RT-qPCR)

Quantification of specific viral RNA was performed using Power SYBR® Green PCR Master Mix (4367659, Applied Biosystems™, Life Technologies Ltd., Woolston Warnington, UK) and QuantStudio3 Real-Time PCR instrument (A28132, Applied Biosystems™). The primer list is shown in Tables 1 and 2.

**Table 1.**
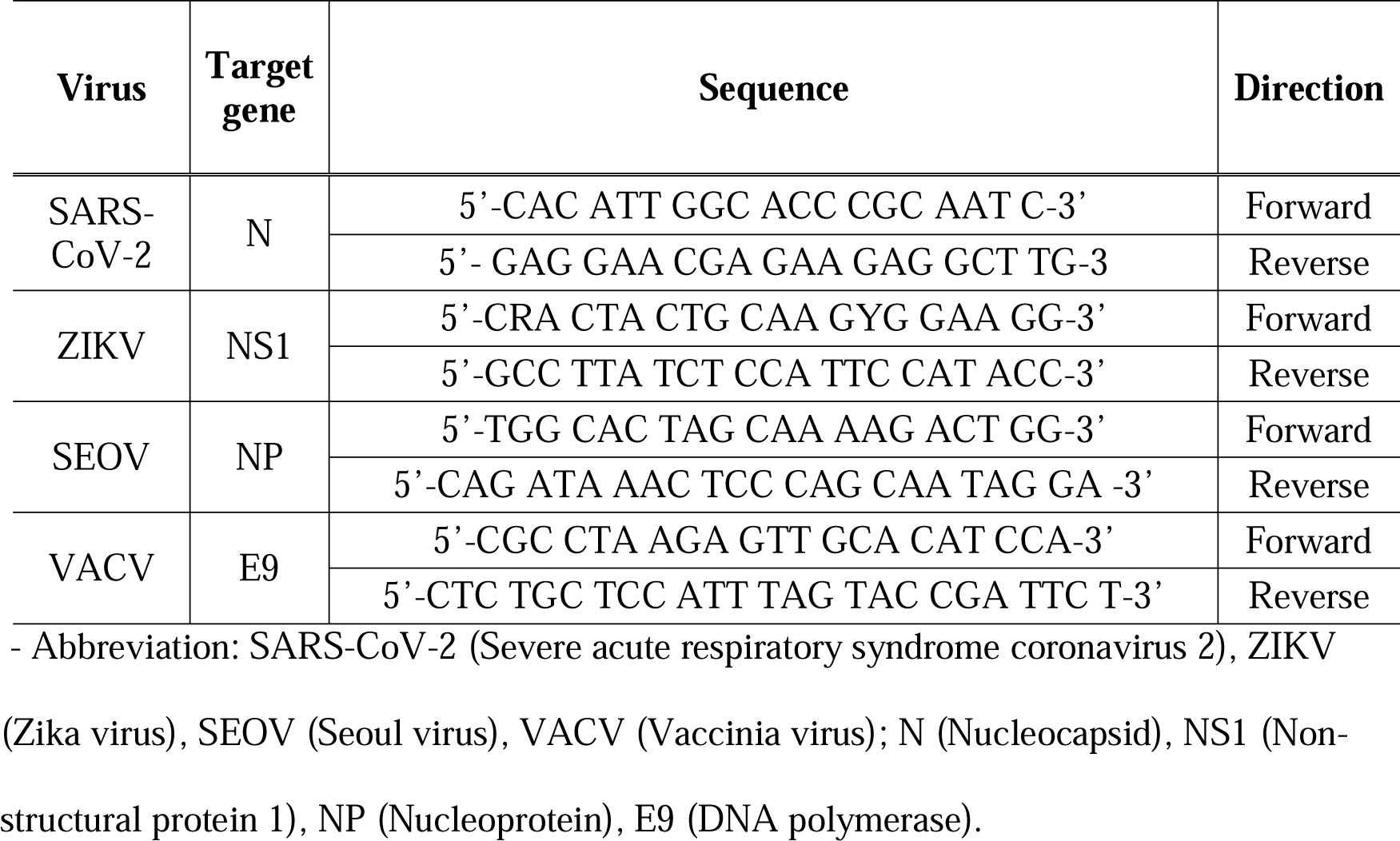
Primer sequences of viruses for RT-qPCR.

**Table 2.**
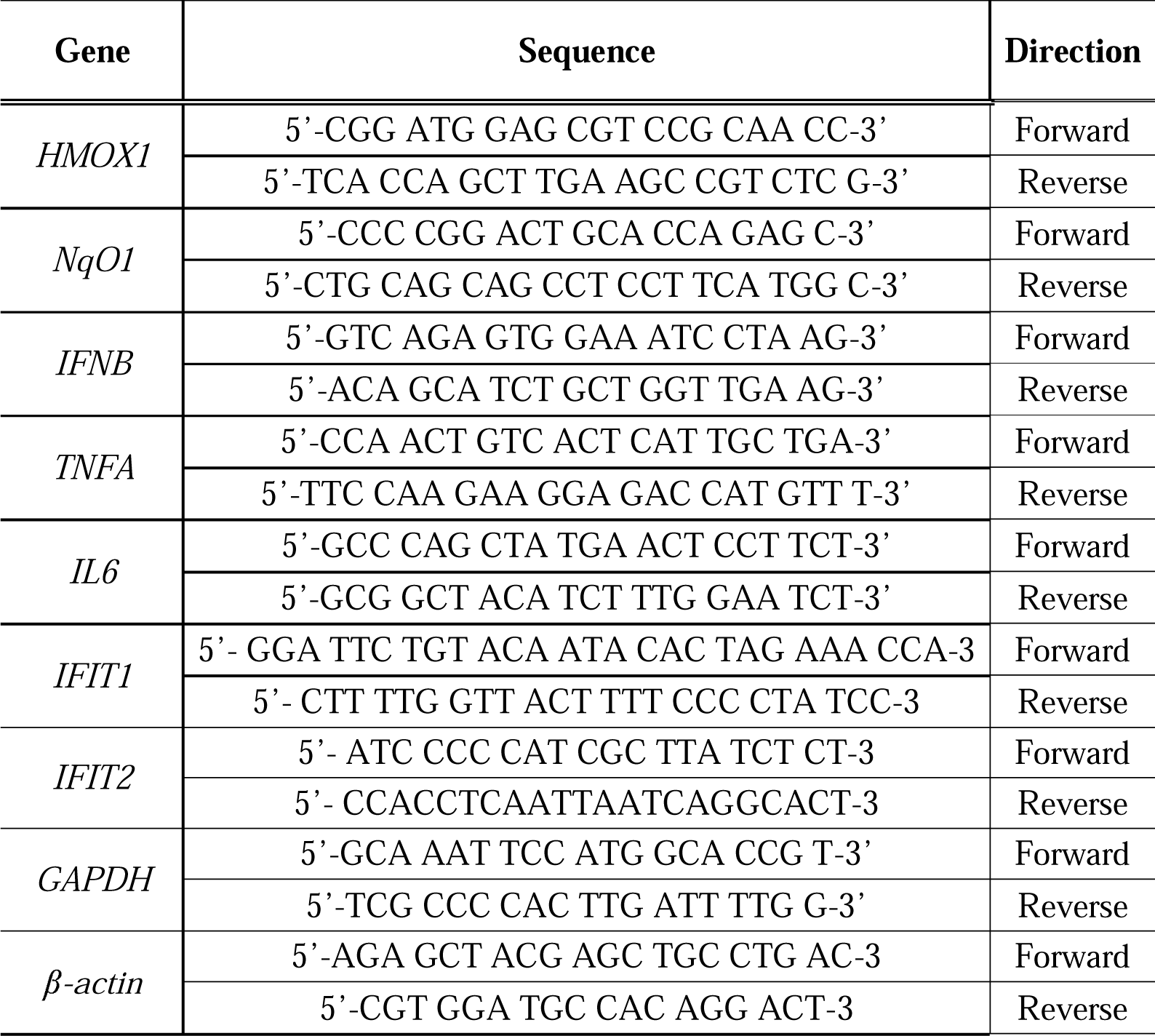
Primer sequences of human-related genes for RT-qPCR.

### 2.7. Plaque assay

Vero E6 cells (1×10^6^ cells per well) were seeded into 6-well plates and incubated overnight at 37°C in 5% CO_2_. Cells were washed with PBS and infected with serial dilutions of SARS-CoV-2 supernatant using serum-free medium. 90 minutes after infection, plates were shaken every 15 minutes for adsorption. Overlay medium (DMEM/F12 medium) containing 0.6% agar was added and incubated at 37°C with 5% CO_2_ for 4 days. Formaldehyde was used for fixation and crystal violet staining was performed.

### 2.8. Western blot

Cell lysis was performed using Radioimmunoprecipitation (RIP) assay buffer (#9806, Cell Signaling Technology, 3 Trask Lane, Danvers, MA 01923, USA) supplemented with protease and phosphatase inhibitor (#5872). SDS-PAGE was utilized to separate the lysed cells, and PVDF membranes 617203, Millipore Ltd. Tullagreen, Carrigtwohill, USA) were used for protein transfer. The membranes were blocked using tris-buffered saline (TBS) and 0.1% Tween-20 (#1706531, Bio-Rad Laboratories, Inc., USA) (TBS-T) with 5% skim milk for 1 h at room temperature (RT). Primary antibodies against SARS-CoV-2 nucleocapsid (PA1-41098, Invitrogen™), SARS-CoV-2 spike protein (S1) (#E5S3V) and GAPDH (G9545, Sigma-Aldrich) were incubated overnight at 4 °C in TBS-T. After three washes with TBS-T, secondary antibodies (111-035-003, ©Jackson ImmunoResearch Inc, West Grove, PA, USA) were applied and incubated for 1 h at RT.

### 2.9. Total RNA-sequencing (Total RNAseq) analysis

TruSeq Stranded Total RNA Library Prep Gold Kit (Illumina, San Diego, CA, USA) was used for library preparation of total RNA sequencing data. The cDNA fragments obtained through RNA sequencing analysis were mapped to the genomic reference (GRCh38) using the HISAT2 program which utilizes the Bowtie2 aligner for spliced read mapping (Kim, Daehwan et al., 2019; Langmead et al., 2012). The processed and mapped reads for each sample were quantified and known genes/transcripts were assembled using the StringTie program with a reference gene model (Pertea et al., 2015). The abundance of transcripts was calculated as read count and normalized using fragments per kilobase of transcript per million mapped reads (FPKM) and transcripts per kilobase million (TPM) values. Differential gene expression analysis (DEG) was performed using the read count values, applying the StringTie-e option for original raw data, filtering genes with low quality, and using the edge R library-calcNormFactors to calculate TMM (Trimmed mean of M-values) normalization (adjusted *p*-value < 0.05; derived from a hypergeometric test & multiple testing correction (FDR) and fold change |Fc| ≥ 2).

### 2.10. Mitochondrial membrane potential staining

The mitochondrial membrane potential probe JC-1 (T3168, Thermo Fisher Scientific) was used to detect the recovery of mitochondrial membrane potential. JC-1 is expressed as a green, fluorescent monomer (∼529 nM) at depolarized and abnormal mitochondrial membrane potentials. In mitochondria with a normally functioning proton pump and normal and hyperpolarized membranes, JC-1 is concentrated inside the mitochondria and forms red fluorescent J aggregates (∼590 nM). To stain hACE2-A549 cells seeded on the cover glass in a 12-well plate, JC-1 was diluted to 10 mM and used at a final concentration of 2 µM in serum free DMEM for 20 min at 37 °C with 5% CO_2_. Control cells were treated with H_2_O_2_ for 20 min. After JC-1 staining, Hoechst 33342 (62249, Thermo Fisher Scientific) was used to stain the nucleus for 5 min at RT. After each staining step, cells were washed three times using pre-warmed PBS and mounted on the slide glass. All steps were carried out with the light blocked.

### 2.11. Statistical analysis

Statistical analyses were performed using GraphPad Prism (Version 8.0.2; GraphPad Software, Inc., La Jolla, CA). The values are presented in the bar graph as the mean ± SD of at least three independent experiments, \**p*□<□0.05, \*\**p* <□0.01, \*\*\**p*□<□0.001, and \*\*\*\**p* < 0.0001 were considered statistically significant.

## 3. Results

### 3.1 MIT-001 exhibits antiviral activity against the B.1 strain of SARS-CoV-2 in vitro

To investigate the antiviral activity of MIT-001 against SARS-CoV-2 B.1, Vero E6 and hACE2-A549 cells were infected at MOI of 0.01 and 0.1, respectively. After 2 h, MIT-001 was administered to infected cells at concentrations of 30, 20, 10, and 1 µM. At 24 hpi, total RNA, protein extract, and supernatant were collected. The dose-dependent effect of MIT-001 on SARS-CoV-2 B.1 Nucleocapsid (N) replication was observed, with a 1000-fold reduction in viral replication at 30 µM concentration (Figure 1A). The EC_50_ value for MIT-001 against SARS-CoV-2 B.1 was 1.77 µM (Figure 1B). Plaque assay analysis using supernatant showed a 1000-fold reduction in infectious virus particles at 30 µM of MIT-001 (Figure 1C), and the expression of N proteins were not detected (Figure 1D). Furthermore, the antiviral activity of MIT-001 against SARS-CoV-2 B.1 in hACE2-A549 cells showed a 100-fold reduction at a concentration of 30 µM (Figure 1E), with an EC_50_ value of 1.46 µM (Figure 1F). The viral particles were reduced by approximately 100-fold (Figure 1G), and N proteins were undetectable at 30 µM treatment in hACE2-A549 cells (Figure 1H).

**Figure 1.**
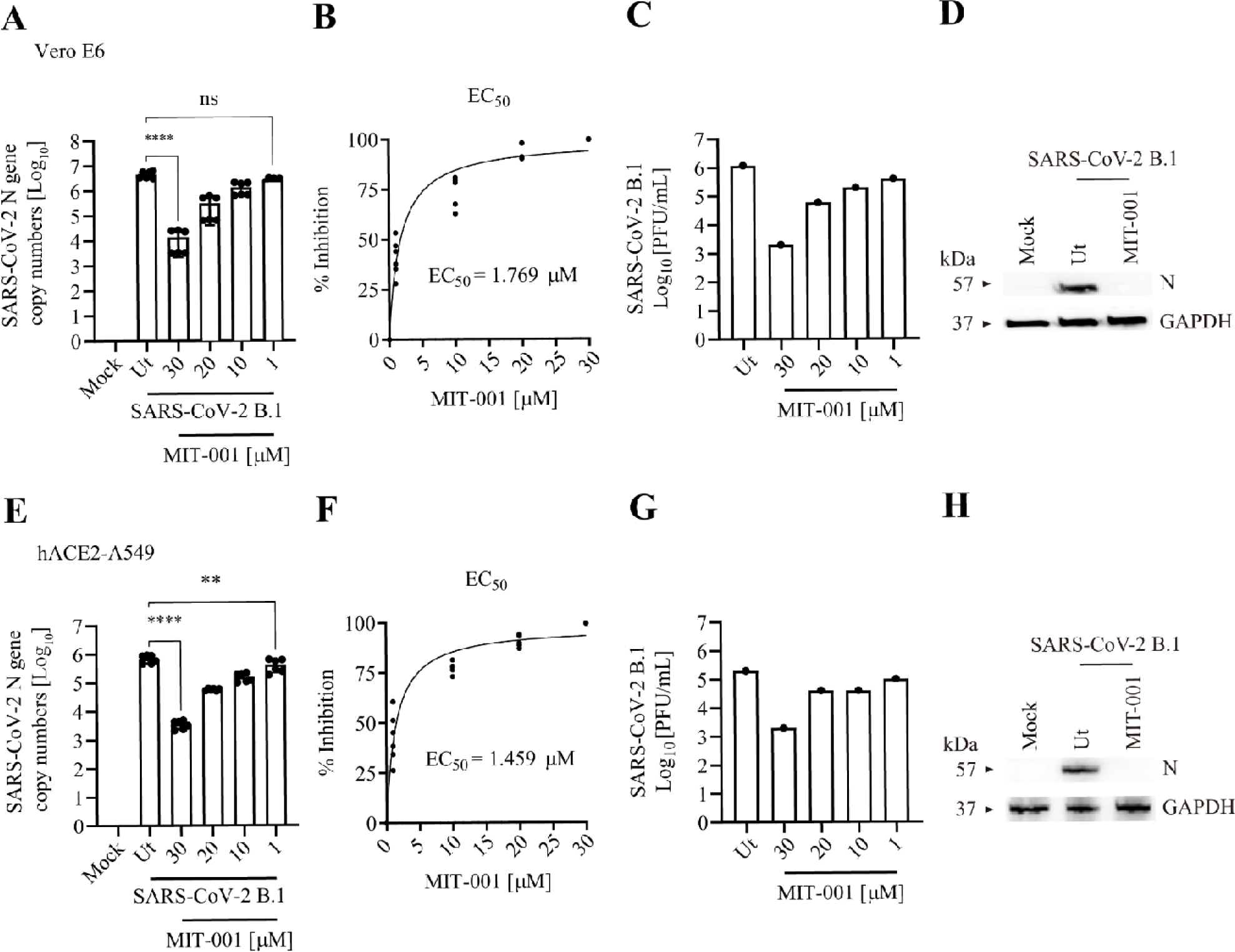
Evaluation of antiviral activity of MIT-001 against the SARS-CoV-2 B.1 strain in Vero E6 and hACE2-A549. Vero E6 and hACE2-A549 cells were infected with SARS-CoV-2 B.1 at MOI=0.01 and 0.1, respectively. 2 h after infection, MIT-001 was treated for each concentration condition and cells and supernatants were harvested 24 hpi. The concentration of MIT-001 processed for Western blot analysis is 30 uM. (A) Analysis of SARS-CoV-2 N gene expression in Vero E6 cells using RT-qPCR. (B) EC_50_ value of MIT-001 in SARS-CoV-2 infected Vero E6 cells. (C) Infectious viral titer determined by plaque assay. (D) SARS-CoV-2 N protein expression in Vero E6 cells after MIT-001 (30 μM) treatment using Western blot. (E) N gene expression in hACE2-A549 cells using RT-qPCR. (F) EC_50_ values. (G) Infectious viral titer in SARS-CoV-2 infected hACE2-A549 cells. (H) Western blot analysis. Data presented are representative of three independent experiments performed in triplicate. \**p*□<□0.05, \*\**p*□<□0.01, \*\*\**p*□<□0.001, and **** *p* < 00001, one-way ANOVA (Ut = Untreated), (ns = non-significant).

### 3.2 MIT-001 inhibits the expression of pro-inflammatory genes but increases the expression of anti-oxidative genes upon SARS-CoV-2 B.1 in hACE2-A549 cells

We evaluated the effect of MIT-001 treatment on inflammatory cytokines and Nrf2-related genes in hACE2-A549 cells infected with SARS-CoV-2 B.1. The total RNAseq analysis showed that SARS-CoV-2 B.1 infection highly induced the interferon (IFN) signaling pathway and pro-inflammatory cytokine genes. However, MIT-001 treatment inhibited the expression of IFN signaling and pro-inflammatory cytokines upon SARS-CoV-2 B.1 infection. (Figure 2A). Regarding Nrf2-related genes, Nrf2 signaling was initially downregulated in SARS-CoV-2 B.1-infected cells; however, MIT-001 treatment led to the upregulation of the Nrf2 pathway. In addition to the upregulation of the heme oxygenase-1 (*HMOX1*) due to MIT-001 treatment, our findings revealed that the restoration of biliverdin (*BLVR*), a heme metabolite, at the RNA level occurred (Figure 2B). RT-qPCR results confirmed the downregulation of *IFNB*, interferon induced protein with tetratricopeptide repeats 1 (*IFIT1)*, *IFIT2*, Interleukin 6 (*IL6)* and tumor necrosis factor alpha (*TNFA)* expression after MIT-001 treatment (Figure 2C). RT-qPCR analysis showed that the expression of *HMOX1* and NAD(P)H: quinone oxidoreductase 1 (*NqO1*), Nrf2-induced genes associated with antioxidant activity, significantly increased after MIT-001 treatment, approaching or surpassing the levels in untreated infected cells (Figure 2D). These results suggest that MIT-001 induces strong anti-inflammatory and anti-oxidant responses in SARS-CoV-2 infected hACE2-A549 cells.

**Figure 2.**
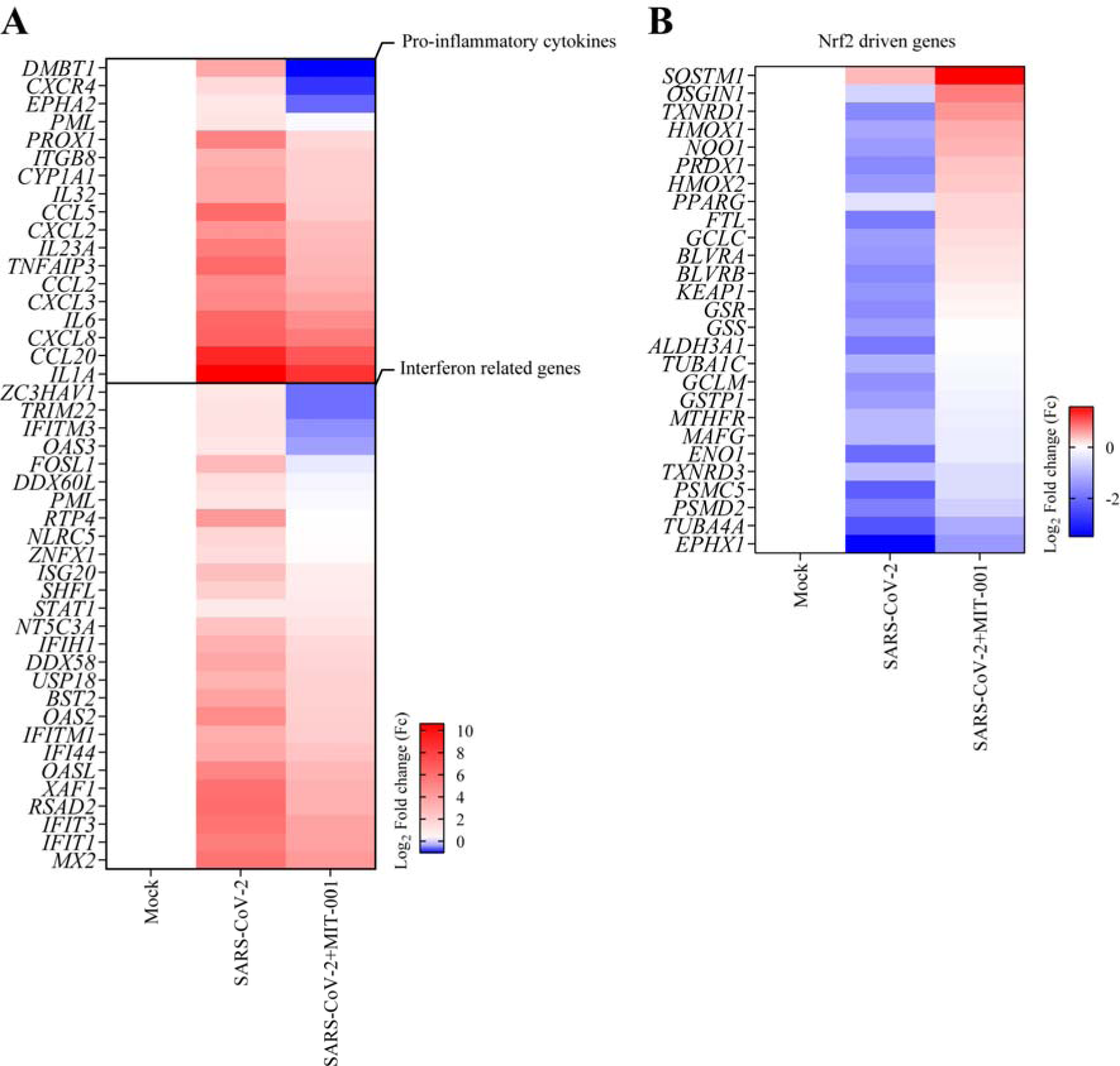
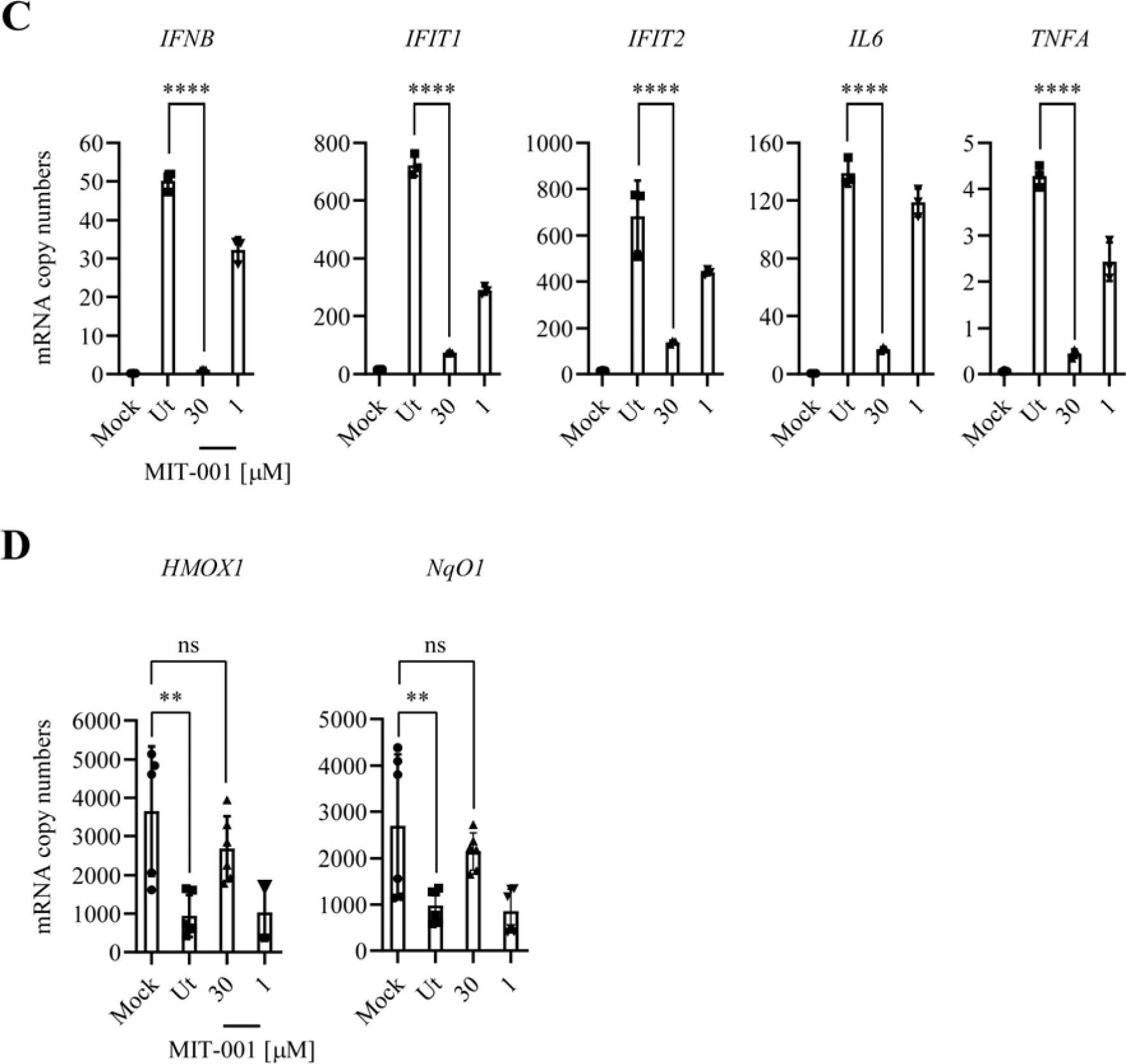
Inflammatory cytokines analysis and up-regulation of Nrf2-related genes after MIT-001 treatment in hACE2-A549 cells infected with the SARS-CoV-2 B.1 strain. hACE2-A549 cells were infected with SARS-CoV-2 B.1 (MOI=0.1). 2h after infection, MIT-001 was treated for 30 and 1 μM. Infected cells were harvested at 24 hpi. (A) Heat map of the inflammation genes expressed in SARS-CoV-2 infected cells after MIT-001 (30 μM) treatment. (B) The expression of Nrf2-related genes in SARS-CoV-2 infected cells after MIT-001 (30 μM) treatment. (C) The expression of interferon genes (*IFNB, IFIT1* and *IFIT2*) and pro-inflammatory cytokines (*TNFA* and *IL6)* in SARS-CoV-2 infected cells after MIT-001 treatment. (D) The expression of Nrf2 related genes (*HMOX1* and *NqO1*) in SARS-CoV-2 infected cells after MIT treatment. Data presented are representative of three independent experiments performed in triplicate. \**p*□<□0.05, \*\**p*□<□0.01, \*\*\**p*□<□0.001, and **** *p* < 0.0001, one-way ANOVA (Ut = Untreated), (ns = non-significant). The total RNAseq data were subjected to a DEG analysis, and the comparative combination satisfied the condition |(Fc)| ≥ 2 & adjust *p*-value < 0.05

### 3.3 MIT-001 rescues mitochondrial membrane potential (ΔΨm) and dynamics of SARS-CoV-2-infected hACE2-A549 cells

To evaluate the homeostasis of mitochondria after MIT-001 treatment, hACE2-A549 cells were infected with the SARS-CoV-2 B.1 at MOI of 0.1. We performed JC-1 staining to detect dynamic changes in the mitochondrial membrane potential (ΔΨm). The ΔΨm (JC-1 aggregate, Red) of hACE2-A549 cells decreased after SARS-CoV-2 infection, and MIT-001 treatment of infected cells resulted in the same or higher intensity as that of the uninfected cells at 24 hpi (Figures 3A and B). This suggests that mitochondria are under constant stress due to SARS-CoV-2 infection and mitochondrial stress is inhibited by MIT-001 treatment.

**Figure 3.**
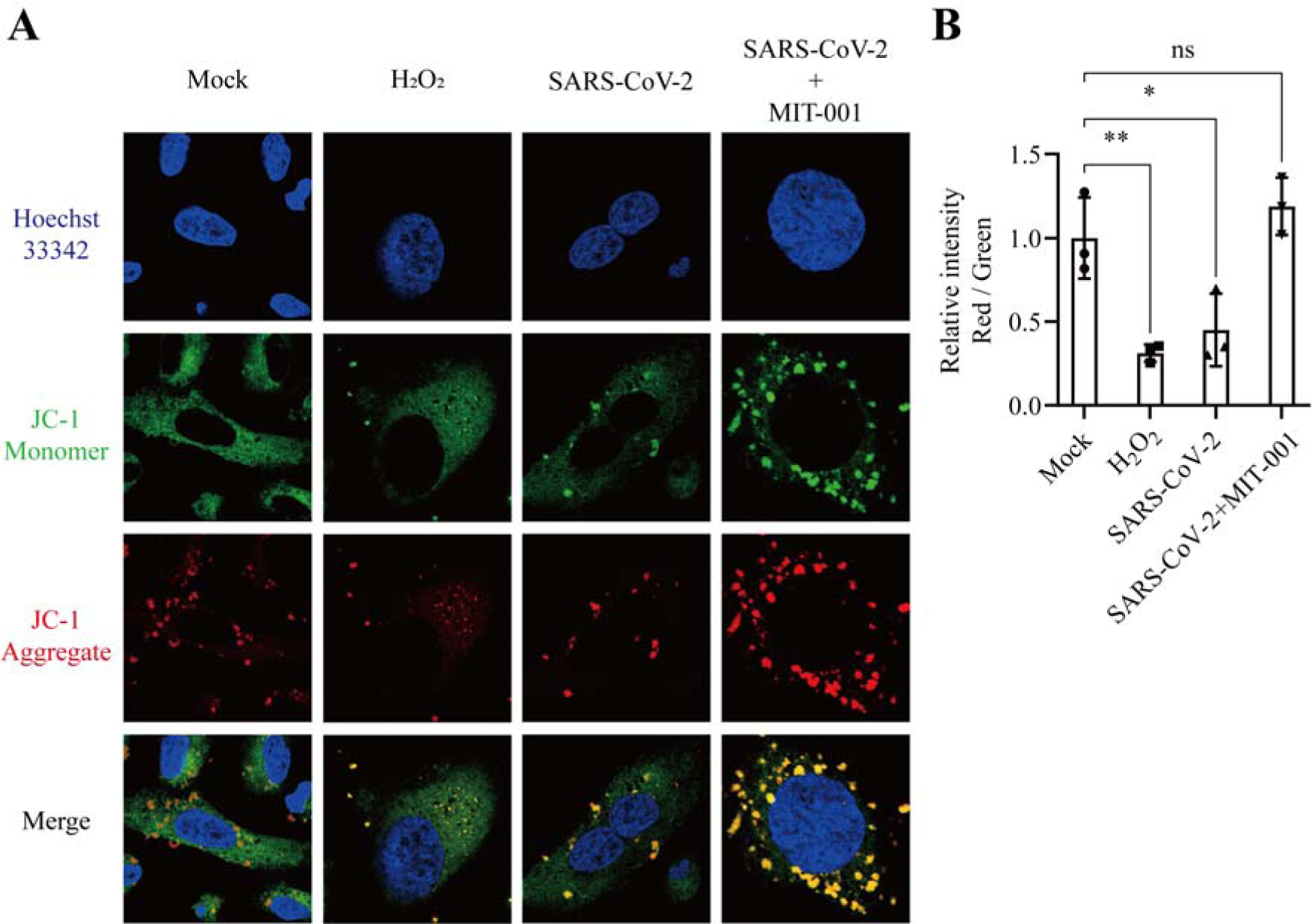
Recovery of mitochondrial homeostasis in SARS-CoV-2 B.1 infected hACE2-A549 after MIT-001 treatment. JC-1 was visible either as green (∼529 nm) for JC-1 monomers or red (∼590 nm) for JC-1 aggregates. Hoecsht33342 (∼488 nm) indicate the nucleus (Blue). Controls were treated with H_2_O_2_ (0.3%). (A) Evaluation of mitochondrial membrane potential using JC-1 staining after MIT-001 (30 μM) treatment in SARS-CoV-2 infected hACE2-A549 cells. (B) The intensity of JC-1 aggregates (Red) / JC-1 monomer (Green) using ImageJ program. Data presented are representative of three independent experiments performed in triplicate. \**p*□<□0.05, \*\**p*□<□0.01, \*\*\**p*□<□0.001, and **** *p* < 0.0001, one-way ANOVA (Ut = Untreated), (ns = non-significant).

### 3.4 MIT-001 exhibits broad-spectrum antiviral activity against SARS-CoV-2 variants and multiple human pathogenic viruses

The antiviral activity of MIT-001 was not restricted to SARS-CoV-2 B.1 but also extended to other human pathogenic viruses. To evaluate broad-spectrum antiviral activity against SARS-CoV-2 variants and multiple viruses, hACE2-A549 and Vero E6 cells were infected with SARS-CoV-2 B.1.617.2 (Delta), BA.1 (Omicron), SEOV, ZIKV, and VACV. Cells were harvested and total RNA was extracted both 24 and 48 hpi. MIT-001 showed approximately 1000-fold and 30-fold antiviral effects against the SARS-CoV-2 B.1.617.2 and BA.1 strains, respectively in hACE2-A549 cells (Figures 4A and B). We observed an approximately 3-fold reduction of SEOV replication in Vero E6 (Figure 4C). Replication of ZIKV was decreased by approximately 16-folds at 48 hpi (Figure 4D). The VACV replication was reduced approximately 15-fold at 30 µM of MIT-001 (Figure 4E). The EC_50_ means according to each antiviral efficacy are summarized in Table 3.

**Figure 4.**
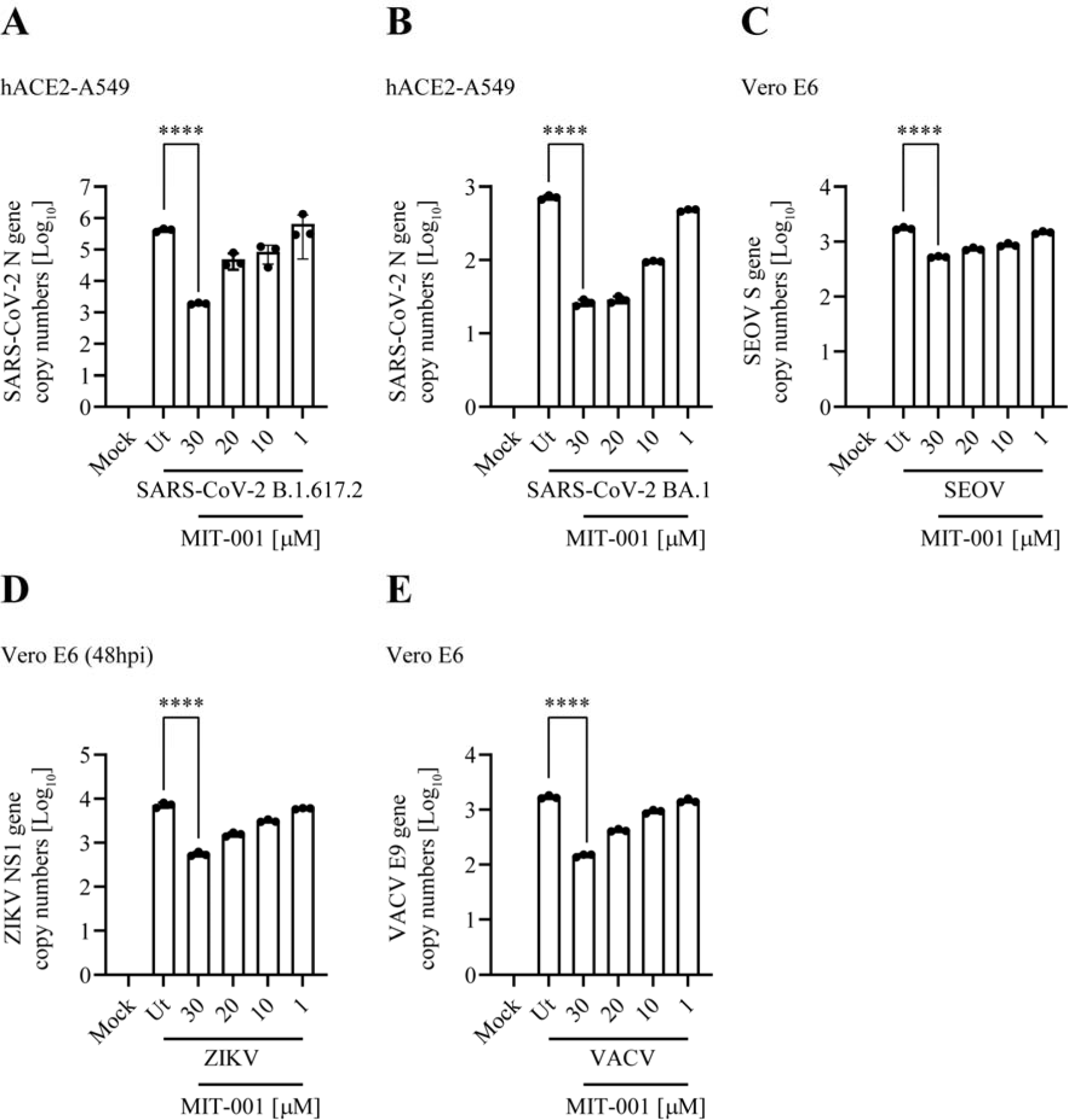
Evaluation of anti-viral activity of MIT-001 against SARS-CoV-2 B.1.617.2, BA.1 and multiple viruses. Cells were infected with SARS-CoV-2 variants, SEOV, ZIKV, and VACV. Total RNA and supernatants were collected at 24 hpi except for ZIKV at 48 hpi. (A) Evaluation of antiviral efficacy against SARS-CoV-2 B.1.617.2 (Delta), N gene expression analysis (hACE2-A549, MOI of 0.1). (B) SARS-CoV-2 BA.1 (Omicron), (hACE2-A549, MOI of 0.1). (C) SEOV, NP gene expression analysis (Vero E6, MOI of 0.01). (D) ZIKV, NS1 gene expression analysis at 48 hpi (Vero E6, MOI of 0.01). (G) VACV, E9 gene expression analysis (Vero E6, MOI of 0.01). Data presented are representative of three independent experiments performed in triplicate. **** *p* < 0.0001, one-way ANOVA (Ut = Untreated).

**Table 3.**
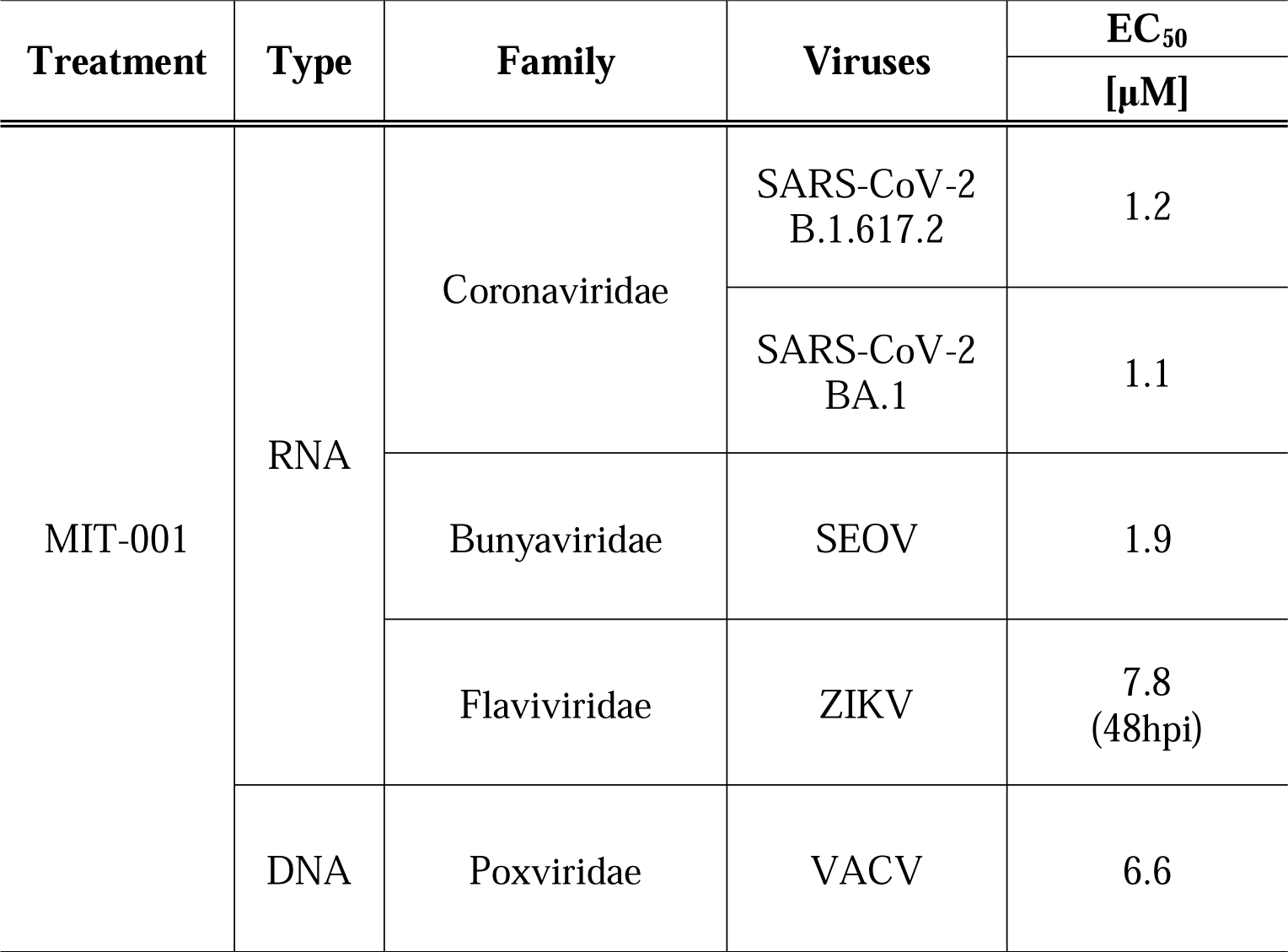
Mean of EC_50_ of MIT-001 against different strains of SARS-CoV-2 and multiple viruses.

| Treatment | Type | Family | Viruses | EC<sub>50</sub> |
| --- | --- | --- | --- | --- |
|  |  |  |  | [μM] |
| MIT-001 | RNA | Coronaviridae | SARS-CoV-2<br>B.1.617.2 | 1.2 |
|  |  |  | SARS-CoV-2<br>BA.1 | 1.1 |
|  |  | Bunyaviridae | SEOV | 1.9 |
|  |  | Flaviviridae | ZIKV | 7.8<br>(48hpi) |
|  | DNA | Poxviridae | VACV | 6.6 |

## 4. Discussion

Emerging SARS-CoV-2 variants and concurrent outbreaks, such as the monkeypox virus, highlight the urgent need for broad-spectrum antiviral therapies (Kemp SA et al., 2021; Minhaj et al., 2022). Mitochondria-targeted broad-spectrum antivirals have shown promise against various viral infections (Gatti et al., 2020). The previous study demonstrated antiviral activity of 4-OI against VACV and ZIKV. Notably, MIT-001 exhibited broad antiviral activity with EC_50_ values ranging from 1.1 to 7.8 µM against SARS-CoV-2 variants (B.1.617.2 and BA.1), SEOV, ZIKV, and VACV. Our findings provide significant evidence that MIT-001, a mitochondria-targeted ROS scavenger, may serve as a novel and comprehensive approach to broad-spectrum antiviral therapeutics.

Viral infections increase ROS activity, leading to oxidative stress and the activation of cytoprotective genes through Nrf2-dependent pathways (Deramaudt et al., 2013; Johnson et al., 2008; Herengt et al., 2021; Osburn et al., 2008; Hayes et al., 2014). However, in recent studies, SARS-CoV-2 infection enabled to inhibit the Nrf2 pathway, as observed in biopsies from COVID-19 patients (Olagnier et al., 2020; Zhang et al., 2022a; Zhang et al., 2022b). The SARS-CoV-2 non-structural protein 14 (nsp14) interacted with Sirtuin 1 (S*IRT1*) to inhibit the activation of Nrf2 pathway (Fratta et al., 2021). The treatment of 4-OI has been shown to activate the Nrf2 pathway in SARS-CoV-2 infected cells by dissociating the Keap1-Nrf2 complex in the cytoplasm. This activation of Nrf2 led to the inhibition of SARS-CoV-2 replication and subsequent pro-inflammatory responses. The transcription factor Nrf2 regulates the expression of antioxidant genes such as *HMOX1* and *NqO1* (Maines et al., 2005; Ma et al., 2013; Tonelli et al., 2018). Previous studies have shown that *HMOX1*-induced *BLVR* mitigated the replication of SARS-CoV-2 and Ebola virus (EBOV) (Olagnier et al. 2020; Fratta et al. 2021; De Clercq et al. 2015). Our study demonstrated that treatment with MIT-001 effectively restored the decreased expression of *HMOX1* caused by SARS-CoV-2 infection to levels comparable to those observed in uninfected cells. Furthermore, the restoration of *HMOX1* gene expression resulted in the up-regulation of *BLVR A* and *B*, which may potentially influence the antiviral activity of MIT-001 against the virus. However, further investigations are required to elucidate the precise mechanism of action underlying the therapeutic effects of MIT-001.

The study has limitations: Firstly, the antiviral activity of MIT-001 was only tested *in vitro*, requiring further evaluation in animal models and humans to assess safety and efficacy. Secondly, a limited number of viruses were tested, necessitating additional research to determine the antiviral activity of MIT-001 against other viruses. Thirdly, the mode of action remains to be further investigated.

In conclusion, MIT-001 provides broad-spectrum antiviral activity against SARS-CoV-2 and multiple zoonotic viruses *in vitro*. This study highlights the potential of MIT-001 for developing a novel mitochondria-targeted antiviral against emerging viral infections.

## Acknowledgments

We thank Dr. Jacob S. Yount (Ohio State University), Seung-Hwan Baek (Hallym University), and Nayeon Jang (Hallym University) for supporting this research.

## Funding sources

This research was supported by Mitoimmune Therapeutics (6R220101510S000100). In addition, this study was provided by Korea Institute of Marine Science & Technology Promotion (KIMST) funded by the Ministry of Oceans and Fisheries, Korea (20210466). It was partially funded by a National Research Foundation of Korea (NRF) grant funded by the Korean government (MSIT) (No. NRF-2021M3H4A4079154).

## Disclosure

The authors declare no competing financial interests.

## Footnotes

1 Abbreviations: COVID-19 - Coronavirus Disease 2019; SARS-CoV-2 – severe acute respiratory syndrome 2 virus;

## Notes

### Competing Interest Statement

The authors have declared no competing interest.

## References

1. World Health Organization., 2020. International Health Regulations Emergency Committee on Novel Coronavirus in China.

2. World Health Organization., 2020. WHO Director-General’s Opening Remarks at the Media Briefing on COVID-19.

3. Saravolatz, L.D., Depcinski, S., Sharma, M., 2023. Molnupiravir and nirmatrelvir-ritonavir: oral coronavirus disease 2019 antiviral drugs. Clin. Infect. Dis. 76, 165–171.

4. Lamb, Y. N., 2022. Nirmatrelvir plus ritonavir: first approval. Drugs, 82(5), 585–591.

5. Olagnier, D., Farahani, E., Thyrsted, J., Blay-Cadanet, J., Herengt, A., Idorn, M., et al., 2020. SARS-CoV2-mediated suppression of NRF2-signaling reveals potent antiviral and anti-inflammatory activity of 4-octyl-itaconate and dimethyl fumarate. Nat. Commun. 11, 4938.

6. Grootaert, M.O., Schrijvers, D.M., Van Spaendonk, H., Breynaert, A., Hermans, N., Van Hoof, V.O., et al., 2016. NecroX-7 reduces necrotic core formation in atherosclerotic plaques of Apoe knockout mice. Atherosclerosis. 252, 166–174.

7. Im, K.-I., Kim, N., Lim, J.-Y., Nam, Y.-S., Lee, E.-S., Kim, E.-J., et al., 2015. The free radical scavenger NecroX-7 attenuates acute graft-versus-host disease via reciprocal regulation of Th1/regulatory T cells and inhibition of HMGB1 release. J. Immunol. 194, 5223–5232.

8. Kim, D., Paggi, J.M., Park, C., Bennett, C., Salzberg, S.L., 2019. Graph-based genome alignment and genotyping with HISAT2 and HISAT-genotype. Nat. Biotechnol. 37, 907–915.

9. Langmead, B., Salzberg, S.L., 2012. Fast gapped-read alignment with Bowtie 2. Nat. Methods. 9, 357–359.

10. Pertea, M., Pertea, G.M., Antonescu, C.M., Chang, T.-C., Mendell, J.T., Salzberg, S.L., 2015. StringTie enables improved reconstruction of a transcriptome from RNA-seq reads. Nat. Biotechnol. 33, 290–295.

11. Kemp, S.A., Collier, D.A., Datir, R.P., Ferreira, I.A., Gayed, S., Jahun, A., et al., 2021. SARS-CoV-2 evolution during treatment of chronic infection. Nat. 592, 277–282.

12. Minhaj, F.S., Ogale, Y.P., Whitehill, F., Schultz, J., Foote, M., Davidson, W., et al., 2022. Monkeypox outbreak—Nine states, May 2022: Weekly/June 10, 2022/71 (23); 764–769.

13. Gatti, P., Ilamathi, H.S., Todkar, K., Germain, M., 2020. Mitochondria targeted viral replication and survival strategies—prospective on SARS-CoV-2. Front. Pharmacol. 11, 578599.

14. Deramaudt, T., Dill, C., Bonay, M., 2013. Regulation of oxidative stress by Nrf2 in the pathophysiology of infectious diseases. Med. Mal. Infect. 43, 100–107.

15. Johnson, J.A., Johnson, D.A., Kraft, A.D., Calkins, M.J., Jakel, R.J., Vargas, M.R., et al., 2008. The Nrf2–ARE pathway: an indicator and modulator of oxidative stress in neurodegeneration. Ann. N. Y. Acad. Sci. 1147, 61–69.

16. Herengt, A., Thyrsted, J., Holm, C.K., 2021. NRF2 in viral infection. Antioxidants. 10, 1491.

17. Osburn, W.O., Kensler, T.W., 2008. Nrf2 signaling: an adaptive response pathway for protection against environmental toxic insults. Mutat. Res. Rev. Mutat. Res. 659, 31–39.

18. Hayes, J.D., Dinkova-Kostova, A.T., 2014. The Nrf2 regulatory network provides an interface between redox and intermediary metabolism. Trends Biochem. Sci. 39, 199–218.

19. Zhang, Q., Bastard, P., Liu, Z., Le Pen, J., Moncada-Velez, M., Chen, J., et al., 2020. Inborn errors of type I IFN immunity in patients with life-threatening COVID-19. Science. 370, eabd4570.

20. Zhang, S., Wang, J., Wang, L., Aliyari, S., Cheng, G., 2022. SARS-CoV-2 virus NSP14 Impairs NRF2/HMOX1 activation by targeting Sirtuin 1. Cell. Mol. Immunol. 19, 872–882.

21. Maines, M.D., 2005. New insights into biliverdin reductase functions: linking heme metabolism to cell signaling. Physiology 20, 382–389.

22. Ma, Q., 2013. Role of nrf2 in oxidative stress and toxicity. Annu. Rev. Pharmacol. Toxicol. 53, 401–426.

23. Tonelli, C., Chio, I.I.C., Tuveson, D.A., 2018. Transcriptional regulation by Nrf2. Antioxid. Redox Signal. 29, 1727–1745.

24. Fratta Pasini, A.M., Stranieri, C., Cominacini, L., Mozzini, C., 2021. Potential role of antioxidant and anti-inflammatory therapies to prevent severe SARS-CoV-2 complications. Antioxidants. 10, 272.

25. De Clercq, E., 2015. Ebola virus (EBOV) infection: therapeutic strategies. Biochem. Pharmacol. 93, 1–10.

26. Mailloux, R.J., Harper, M.-E., 2012. Mitochondrial proticity and ROS signaling: lessons from the uncoupling proteins. Trends Endocrinol. Metab. 23, 451–458.

27. Sukumar, M., Liu, J., Mehta, G.U., Patel, S.J., Roychoudhuri, R., Crompton, J.G., et al., 2016. Mitochondrial membrane potential identifies cells with enhanced stemness for cellular therapy. Cell Metab. 23, 63–76.

28. Santel, A., Fuller, M.T., 2001. Control of mitochondrial morphology by a human mitofusin. J. Cell Sci. 114, 867–874.

29. Jahani, M., Dokaneheifard, S., Mansouri, K., 2020. Hypoxia: A key feature of COVID-19 launching activation of HIF-1 and cytokine storm. J. Inflamm. 17, 1–10.

30. Danta, C.C., 2021. SARS-CoV-2, hypoxia, and calcium signaling: the consequences and therapeutic options. ACS Pharmacol. Transl. Sci. 4, 400–402.

31. Duan, X., Tang, X., Nair, M.S., Zhang, T., Qiu, Y., Zhang, W., et al., 2021. An airway organoid-based screen identifies a role for the HIF1α-glycolysis axis in SARS-CoV-2 infection. Cell Rep. 37.

32. Díaz-Resendiz, K.J.G., Covantes-Rosales, C.E., Benítez-Trinidad, A.B., Navidad-Murrieta, M.S., Razura-Carmona, F.F., Carrillo-Cruz, C.D., et al., 2022. Effect of fucoidan on the mitochondrial membrane potential (ΔΨm) of leukocytes from patients with active COVID-19 and subjects that recovered from SARS-CoV-2 infection. Mar. Drugs 20, 99.

